# Complex interplay of biomechanics and ecology influenced crab claw morphology evolution

**DOI:** 10.64898/2026.06.23.733945

**Authors:** Russell D. C. Bicknell, Joanna M. Wolfe, John J. Flynn, Adiël A. Klompmaker, Morgan Chase, Phoebe Fu, Melanie J. Hopkins

## Abstract

True crabs (Brachyura) are among the most iconic marine arthropods, representing noteworthy examples of morphological and ecological disparity. A striking feature of brachyurans are their anterior pincer-like appendages: chelipeds. These structures showcase a large diversity of morphologies that reflect ecology and overall multifunctionality. Yet, a comprehensive assessment of appendage functional morphology within phylogenetic and ecological trait contexts has never been attempted. By combining 3D geometric morphometrics, finite element analyses, multilocus molecular phylogeny, and ecological trait data for 80 crab species, including three fossil forms, we unveil a complex evolutionary history for crab chelipeds. Despite extreme shape diversity amongst chelipeds, stress distributions are very similar across taxa and hint a many-to-one pattern. High concentrations of chelipeds within constrained morphospace regions associated with peak pinch forces illustrates that brachyuran morphologies optimised for shell crushing may have arisen in the Cretaceous. Deviations from this morphospace highlight the diversification of non-shell-crushing life modes and the influence of sexual selection on appendages. Neither cheliped shape nor pinch force show phylogenetic signal. Together these results indicate that the evolution of cheliped shape is closely associated with, and inferred to have been strongly influenced by, crab ecology, biomechanical needs and sexual selection.

**SIGNIFICANCE STATEMENT:** Chelipeds, the pincer-like claws of crabs, are among the most morphologically diverse appendages within Arthropoda, yet the evolutionary forces driving this diversity remain poorly understood. By integrating 3D geometric morphometrics, biomechanical modelling, molecular phylogeny, and ecological data across 80 crab species including fossil forms, we demonstrate that cheliped morphology is driven by ecology, biomechanical demands, and sexual selection rather than phylogenetic relatedness. The multifunctionality of these structures produces strong evidence for many-to-one mapping of form to function.

Morphologies optimised for durophagy appear to have originated in the Cretaceous, with subsequent diversification into manipulative and sexually selected forms from a morphologically flexible foundation. These findings demonstrate that cheliped diversity reflects a complex interplay between ecological specialisation, biomechanical optimisation, and sexual selection across Brachyura.

## INTRODUCTION

Arthropods are the most diverse and abundant animal group on Earth (1, 2). Their modular body plan, segmented appendages, and highly modifiable exoskeletons are core aspects of their success (3–6). The diverse array of shapes, sizes, and life modes has made arthropods a fundamental group for exploring evolutionary questions (2, 5, 7–10). They therefore offer an ideal means of examining morphological innovation and ecological diversification.

Brachyuran decapod crustaceans are one of the most well-known modern and fossil marine arthropod groups (11–16). Their claws (chelipeds) show remarkable morphological diversity (13, 17–20) that reflect the multiple appendage functions (e.g., feeding, attracting mates) (12, 19–22). Chelipeds are commonly used for predation and can produce higher reaction forces relative to body-mass than in other animals (23). As such, some brachyurans have been primary predators of shelled molluscs (21, 24, 25) using mechanically simple lever structures that inversely relate force output to mobility and speed (11, 26, 27). Brachyuran chelipeds also often show heterochely—the condition where claws are asymmetrical, differing in morphology, size, or both, reflecting the varied roles of each appendage (17, 18, 28). Chelipeds thus are mechanically simple structures, ideal for understanding links between shape, function, and ecology, that also show substantial morphological complexity (19, 29).

To date, cheliped form among brachyurans has been studied primarily using 2D geometric morphometrics. While this method can distinguish appendages of different taxa (17, 18, 30, 31), fixed landmarks have limited power to separate propodal sections at the species level (30). There has been some use of 3D data for spherical harmonic analyses of Paguroidea appendages (32), but these approaches have not modelled propodus internal regions. A substantial gap in crustacean biology and morphology therefore remains to be explored (22, 33) and chelipeds are a focal point for understanding crab ecology and behaviour (34).

Here we build complete, detailed 3D models of chelipeds from 80 brachyuran species, including three fossil species, spanning 20 superfamilies encompassing a diverse range of appendage shapes (Figure 1). We use these models to estimate the biomechanics and functional performances of these structures and assess how performance is distributed across diverse cheliped morphologies. We then compare functional performance within phylogenetic and morphological frameworks to address how ecological variables (e.g., diet, habitat medium, and signalling behaviour) influenced cheliped evolution.

**Figure 1:**
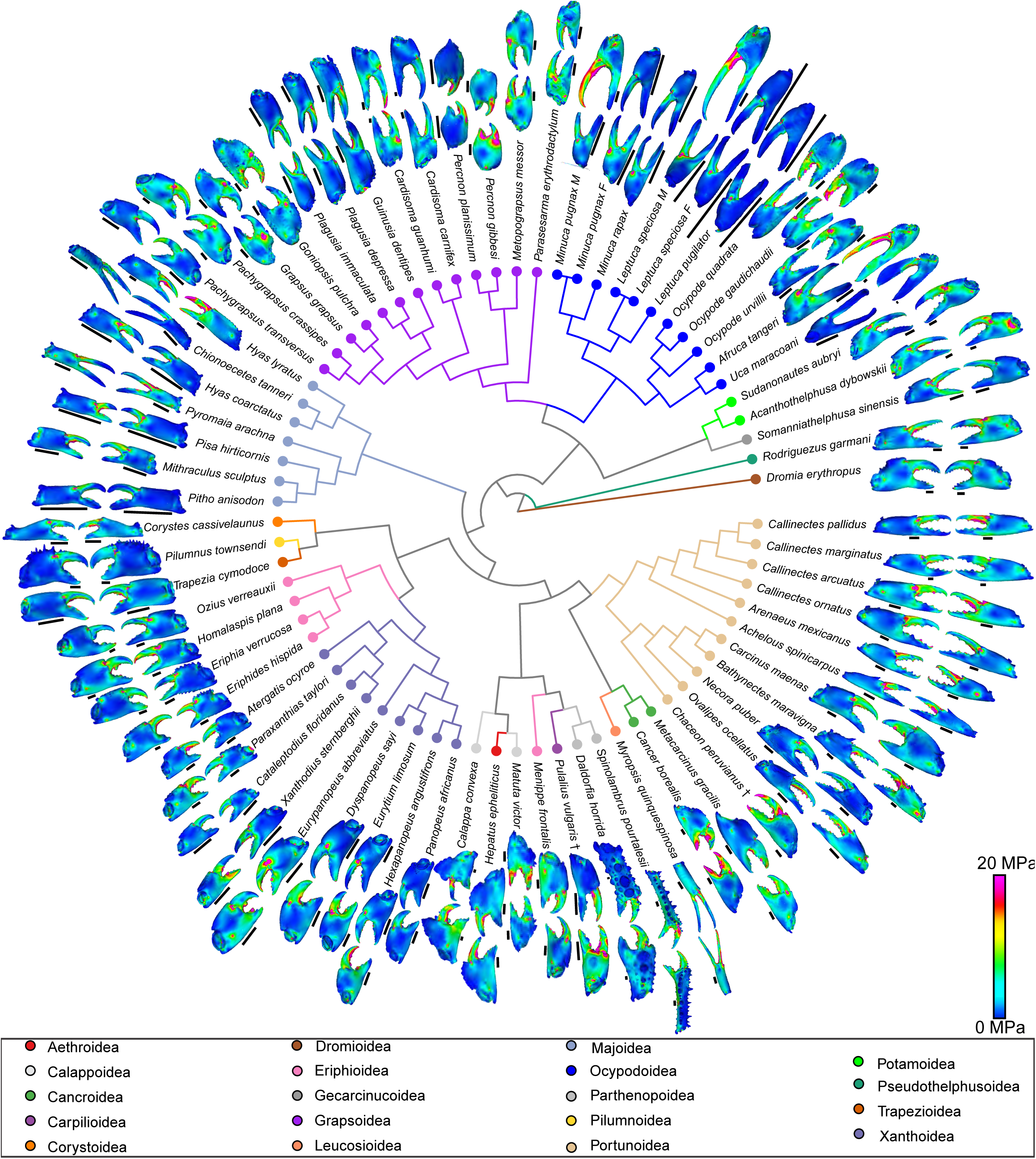
**Molecular phylogeny and 3D finite element models**. Braches and terminals colour coded for superfamilies. Left chelipeds are the inner models, right chelipeds are outer models. Where appropriate, appendages were mirrored. Branches are not scaled. Scale bars: all 5 mm. † are extinct forms.

## RESULTS

### Phylogeny

The phylogenetic results (Figure 1; details in Supplemental Figures 3–5) are mostly consistent regardless of optimality criterion. Inclusion of confamilial molecular data to represent fossils (*Chaceon peruvianus* and *Pulalius vulgaris*) does not significantly impact topology or support values (Supplemental Figures 3, 4).

The topology broadly corresponds to that in Wolfe*, et al.* (14), reflecting utility of comparable data herein. As in several recent studies (14, 35, 36), Heterotremata is paraphyletic and contains a monophyletic Thoracotremata. In the ML analysis, Asian and African freshwater crabs (Gecarcinucoidea and Potamoidea) form the sister group to Thoracotremata with weak support (UFboot = 52). Unlike most previous results, an American freshwater group, Pseudothelphusoidea, forms the sister group to all other Heterotremata (also with weak support: UFboot = 78, pp = 0.9).

### Phylomorphospaces

***Left appendages***: The first principal coordinate (PCO1) for the left appendage describes 41.7% of variation (Figure 2A). Along this axis, morphology ranges from slender chelipeds with elongated fingers (-PCO1) to more rounded, stouter chelipeds, with shorter finger lengths (+PCO1). The PCO2 axis describes 6.9% of variation. Along this axis, chelipeds range from rounded (-PCO2) to more triangular manus morphologies with smaller pollices (+PCO2).

**Figure 2:**
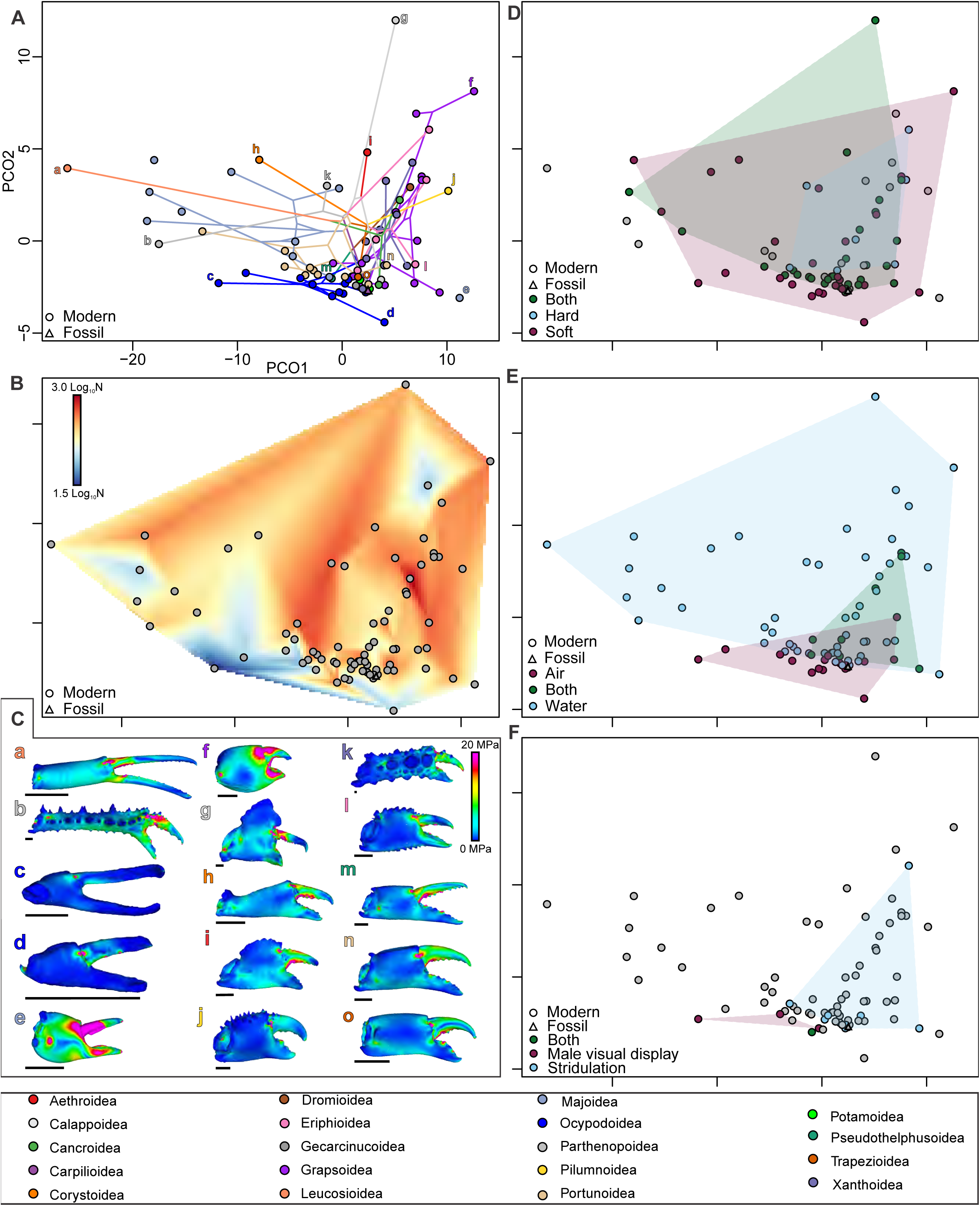
Phylomorphospace, functional landscape, trait space, and finite element models for left chelipeds. (A) Phylomorphospace with branches and terminals are coded for superfamily. Lower case letters correspond to FEMs in (C). Modern and fossil forms are presented in distinct shapes. (B) Functional landscape of NRFs. Modern and fossil forms are presented in distinct shapes. (C) Representative FEMs. Letters colour coded for superfamily. (a) *Myropsis quinquespinosa*. (b) *Spinolambrus pourtalesii*. (c) *Uca maracoani*, male. (d) *Leptuca pugilator*, female. (e) *Paradasygyius*L*depressus*. (f) *Percnon gibbesi*. (g) *Calappa convexa.* (h) *Corystes cassivelaunus.* (i) *Hepatus epheliticus*. (j) *Pilumnus townsendi*. (k) *Daldorfia horrida*. (l) *Eriphia verrucosa*. (m) *Rodriguezus garmani*. (n) *Carcinus maenas*. (o) *Trapezia cymodoce*. All scale bars: 5 mm. (D) Morphospace mapped with diet traits. Grey points are species lacking any data. Modern and fossil forms are presented in distinct shapes. (E) Morphospace mapped with habitat traits. Grey points are species lacking any data. Modern and fossil forms are presented in distinct shapes. (F) Morphospace mapped with sexual display traits. Grey points are species lacking any data, or are female fiddler crabs that lack sexual display. Modern and fossil forms are presented in distinct shapes. Axes in (B, D, E, F) are the same as (A), so are not reproduced.

The phylomorphospace illustrates that Leucosioidea, Majoidea, and Parthenopoidea are sparsely distributed in -PCO1 space, with Eriphioidea, Grapsoidea, Pilumnoidea, and Xanthoidea occupying more of the +PCO1 space. The highest concentration of other superfamilies is about the PCO1 origin (∼0). Notably, Ocypodoidea has a slightly broader distribution towards the morphospace margins. There is no phylogenetic signal in left cheliped shape (*Kmult* = 0.0009, *p* = 0.59).

### Right appendages

PCO1 for the right appendage describes 49.4% of variation (Figure 3A). Along this axis, morphology ranges from slender chelipeds with elongated fingers and manus sections (-PCO1) to round and shorter chelipeds with shorter finger lengths (+PCO1). PCO2 describes 5.7% of variation. Along this axis, cheliped morphologies range from smaller manus sections with elongated fingers (-PCO2) to elongated manus sections with shorter fingers (+PCO2).

**Figure 3:**
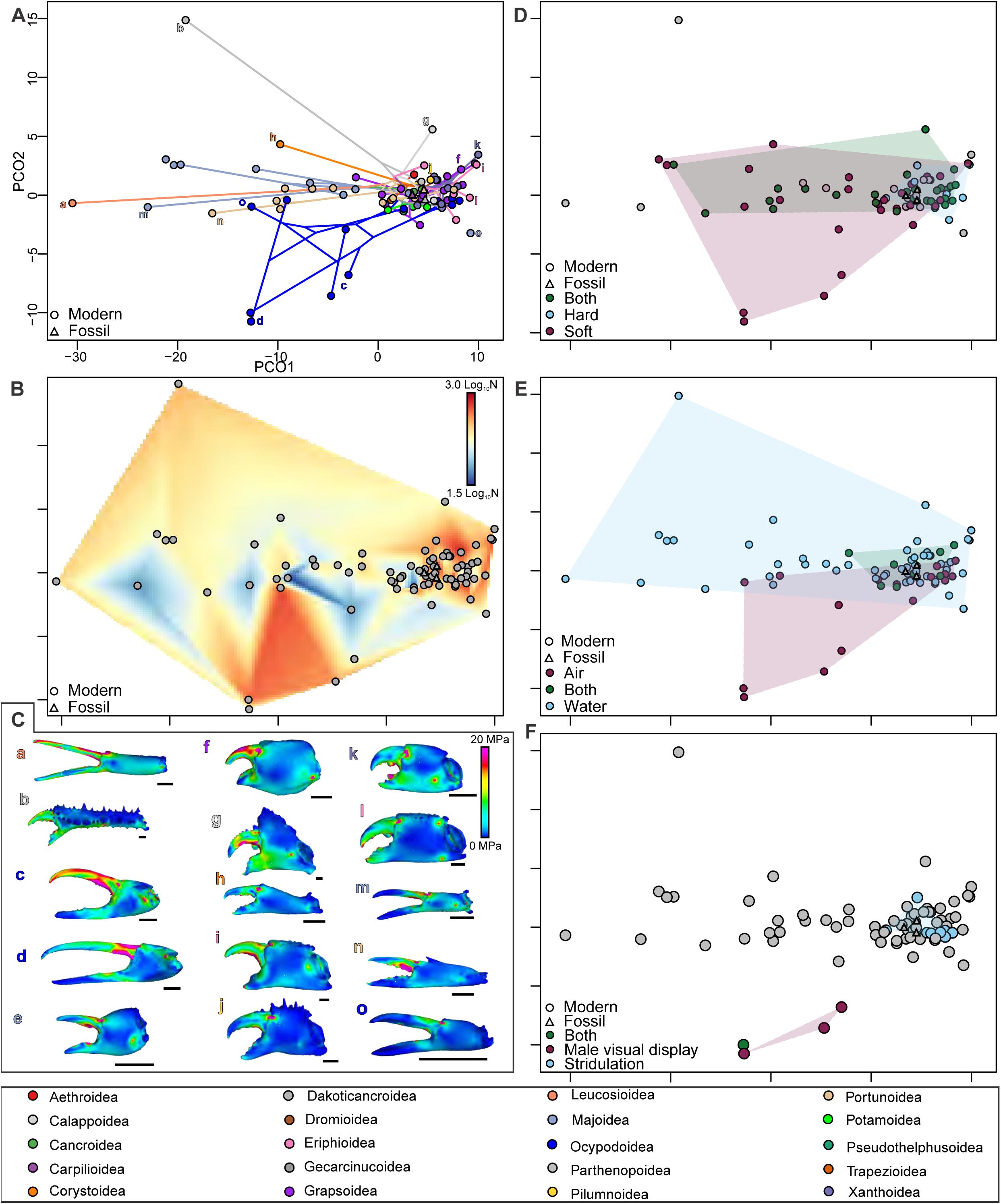
Phylomorphospace, functional landscape, trait space, and finite element models for right chelipeds. (A) Phylomorphospace with branches and terminals are coded for superfamily. Lower case letters correspond to FEMs in (C). Modern and fossil forms are presented in distinct shapes. (B) Functional landscape of NRFs. Grey points represent models. Modern and fossil forms are presented in distinct shapes. (C) Representative FEMs. Letters colour coded for superfamily. (a) *Myropsis quinquespinosa*. (b) *Spinolambrus pourtalesii.* (c) *Minuca pugnax*, male. (d) *Leptuca speciosa*, male. (e) *Paradasygyius*L*depressus*. (f) *Percnon gibbesi*. (g) *Calappa convexa*. (h) *Corystes cassivelaunus*. (i) *Eriphides hispida*. (j) *Pilumnus townsendi*. (k) *Eurypanopeus abbreviatus*. (l) *Eriphia verrucosa*. (m) *Pyromaia arachna*. (n) *Achelous spinicarpus*. (o) *Minuca rapax*. All scale bars: 5 mm. (D) Morphospace mapped with diet traits. Grey points are species lacking any data. Modern and fossil forms are presented in distinct shapes. (E) Morphospace mapped with habitat traits. Grey points are species lacking any data. Modern and fossil forms are presented in distinct shapes. (F) Morphospace mapped with sexual display traits. Grey points are species lacking any data. Modern and fossil forms are presented in distinct shapes. Axes in (B, D, E, F) are the same as (A), so are not reproduced.

The phylomorphospace illustrates a sparse distribution of Leucosioidea, Majoidea, and Parthenopoidea in -PCO1 space, contrasted with a cluster of nearly all other superfamilies in +PCO1 space. About the PCO1 origin (0) Ocypodoidea and Portunoidea dominate, with additional representation from Corystoidea and Potamoidea. There is no phylogenetic signal for right cheliped shape (*Kmult* = 0.001, *p* = 0.50).

### Both appendages

PCO1 for the combined appendage dataset describes 47.9% of variation (Figure 4A). Along this axis, morphologies range from slender chelipeds with elongated manus sections and elongated fingers (-PCO1) to more rounded and shorter chelipeds with shorter fingers (+PCO1). PCO2 describes 5.3% of variation. Along this axis, morphologies range from chelipeds with smaller manus sections and elongated fingers (-PCO2) to rounded manus sections with shorter fingers (+PCO2).

**Figure 4:**
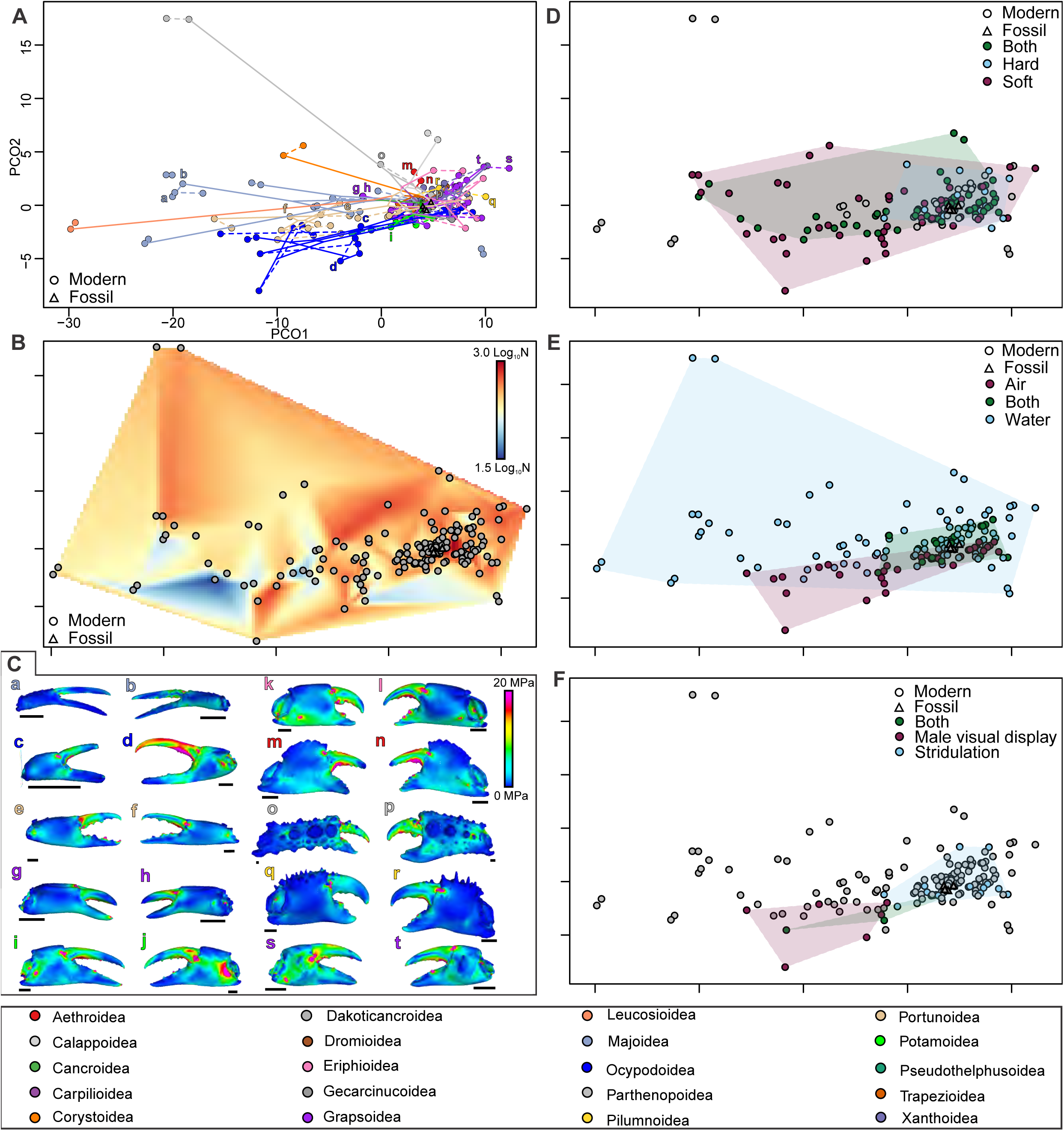
Phylomorphospace, functional landscape, trait space, and finite element models for both appendages. (A) Phylomorphospace with branches and terminals are coded for superfamily. Dotted lines join models of the same specimen. Lower case letters correspond to FEMs in (C). Modern and fossil forms are presented in distinct shapes. (B) Functional landscape of NRFs. Grey points represent models. (C) Representative FEMs. Letters colour coded for superfamily. (a, b) *Chionoecetes tanneri*. (a) Left cheliped. (b) Right cheliped. (c, d) *Minuca pugnax*. (c) Left cheliped. (d) Right cheliped. (e, f) *Callinectes ornatus*. (e) Left cheliped. (f) Right cheliped. (g, h) *Plagusia depressa*. (g) Left cheliped. (h) Right cheliped. (i, j) *Acanthothelphusa dybowskii*. (i) Left cheliped. (j) Right cheliped. (k, l) *Menippe frontalis*. (k) Left cheliped. (l) Right cheliped. (m, n) *Hepatus epheliticus*. (m) Left cheliped. (n) Right cheliped. (o, p) *Daldorfia horrida*. (o) Left cheliped. (p) Right cheliped. (q, r) *Pilumnus townsendi*. (q) Left cheliped. (r) Right cheliped. (s, t) *Parasesarma erythrodactylum*. (s) Left cheliped. (t) Right cheliped. (D) Morphospace mapped with diet traits. Grey points are species lacking any data. Modern and fossil forms are presented in distinct shapes. (E) Morphospace mapped with habitat traits. Grey points are species lacking any data. Modern and fossil forms are presented in distinct shapes. (F) Morphospace mapped with sexual display traits. Grey points are species lacking any data. Modern and fossil forms are presented in distinct shapes. Axes in (B, D, E, F) are the same as (A), so are not reproduced.

The phylomorphospace mirrors the distribution pattern of the right appendage alone. Leucosioidea, Majoidea, and Parthenopoidea are sparsely distributed within -PCO1 space, and most other superfamilies are clustered in +PCO1 space. About the PCO1 origin (0) Ocypodoidea and Portunoidea occur along with members of Corystoidea and Potamoidea, as in the right appendage analysis.

### Finite Element Models

Stress distributions for chelipeds were generally consistent across all models—higher stresses along the pollex and dactylus and lower VM stress within the manus (Figures 1, 2C, 3C, 4C). The highest VM stresses were located about the propodus-dactylus articulation and at constrained denticles (teeth). Taxa showing little to no heterochely (e.g., *Achelous*, *Callinectes*) have similar VM stress distributions, while those with substantial heterochely (e.g., *Daldorfia horrida*, male *Minuca pugnax*, male *Uca maracoani*) show the smaller (often left) cheliped with lower VM stress and the larger (often right) cheliped with higher VM stress. Stouter appendages with pronounced denticles have lower VM stresses within the manus (e.g., *Cataleptodius floridanus, Eriphia verrucosa*, *Pilumnus townsendi*). Forms with higher VM stresses along the fingers also have elongated pollices and dactyli (e.g., *Corystes cassivelaunus*, male *Leptuca speciosa,* male *Minuca pugnax*, *Myropsis quinquespinosa*, and *Pyromaia arachna*). Chelipeds of female fiddler crabs and left appendages of male fiddler crabs (*Minuca pugnax*, *Uca maracoani, Leptuca pugilator*) all show lower VM stresses across the models on average. Notably, higher VM stress distributions across the entire model were observed in *Afruca tangeri*, *Cancer borealis*, *Parasesarma erythrodactylum*, and *Percnon gibbesi*.

### Functional landscape

#### Left appendages

Gradations between lower and higher NRFs are observed along PCO2 (Figure 2B). Morphologies with especially negative PCO2 values, and particularly in the lower left quadrant, generally show a lower NRF. One distinct region of lower NRF also occurs in the upper right quadrant, at about +PCO1 (5) and +PCO2 (8). Most morphologies along +PCO1 and within +PCO2 space show higher NRFs. There is no phylogenetic signal for NRF of left appendages (*K* = 0.006, *p* = 0.12).

#### Right appendages

Gradations between lower and higher NRFs occur in the morphospace across PCO1 and PCO2 (Figure 3B). Peaks in NRFs occur in the right central part of the morphospace (highly +PCO1 and mean PCO2 values) and in the lower central part of the morphospace (moderately -PCO1 to very -PCO2 values). Morphologies about the PCO2 origin (0) along -PCO1 show generally lower NRFs. There is no phylogenetic signal for NRF of right appendages (*K* = 0.0002, *p* = 0.95).

#### Both appendages

Gradations between lower and higher NRFs are present across this landscape (Figure 4B). Morphologies in +PCO2 (>-3) space along PCO1 show higher NRFs. Notable peaks of NRF occur at higher PCO1 values (>-5) on the righthand side of the morphospace. Areas of lower NRFs occur in -PCO2 space along much of the PCO1 axes.

### Trait spaces

#### Left appendages

Brachyurans with mixed (“both”) and soft-bodied diets are spread across morphospace, but durophagous (“hard”) forms are constrained to a small region around +PCO1 and mean PCO2 values (Figure 2D). Aquatic (“water”) forms occur across all parts of PCO1 and PCO2 space, whereas forms with a terrestrial or semi-terrestrial lifestyle (“air”) occur only in -PCO2 spaces and in a narrower range of PCO1 space, and taxa that live in both environments cluster in only a small part of the morphospace (near mean PCO1 and mean PCO2) (Figure 2E). Forms that employ stridulation are spread across higher PCO1(>-3) space, while those that use chelipeds in male visual display, or in both stridulation and display, are restricted to the lower central part of the space [-PCO2 (<0) and moderately - PCO1 (<0)] (Figure 2F). Those using chelipeds only in stridulation or only in male visual display do not overlap in PCO space.

#### Right appendages

For right appendages, brachyurans with mixed and soft-bodied diets are spread across PCO1, while durophagous forms are restricted to very high PCO1 values (>0) (Figure 3D). Taxa with soft diets have a distribution in more negative PCO2 space (-10–+4) compared with species with durophagous or mixed diets. Aquatic forms are present along the entire upper part of the morphospace, generally at the mean or higher values of PCO2, whereas species living only on land occupy the lower center to right portion of the morphospace, and forms that occupy both environments are restricted to a small area only in the central right region (high PCO1 and near mean PCO2 values) (Figure 3E). Forms that use stridulation cluster tightly in only +PCO1 (>1) space, while those that use visual displays, or both stridulation and display, are located in the lower and central regions of the space (-PCO2 and moderately negative PCO1 values), below all other taxa in PCO2 values (Figure 3F).

#### Both appendages

Brachyurans with mixed and soft-bodied diets overlap in the mid to lower parts of the PCO2 morphospace, ranging widely across PCO1 axes, whereas durophagous forms are concentrated in the right central region (high PCO1 values and mean PCO2 values) (Figure 4D). Aquatic forms are present along the entire PCO1 and PCO2 space, whereas species living only on land cluster tightly in -PCO2 (<0) space ranging across only mid- to high positive values of PCO1 (with none in highly negative PCO1 space), while forms that occupy both environments cluster tightly at high PCO1 (>-2) and near mean PCO2 values (Figure 4E). Forms that use stridulation alone cluster tightly at high PCO1 (>-1) and near mean PCO2 values, while those that use male visual displays only are located together in -PCO1 (<-2) and -PCO2 (<0) space, and those that use both stridulation and display span the regions occupied by the other two categories (Figure 4F).

### Fossil forms

In PCO morphospaces for left, right and combined chelipeds, extinct species cluster in +PCO1 (>0) and -PCO2 (<0) space (Figures 2D–F, 3D–F, 4D–F). In dietary plots, *Pulalius vulgaris* (Lincoln Creek Formation, Washington State, USA, Eocene), the only fossil with a left appendage preserved, falls within those species that have a mixed hard and soft diet. For the right appendages, *Dakoticancer australis* (Coon Creek Member, Ripley Formation, Mississippi, USA. Cretaceous [early Maastrichtian]), *Chaceon peruvianus* (no formation data, Patagonia, Pleistocene) and *Pulalius vulgaris* cluster closest to forms with both diets (or unknown) and within the region of morphospace that contains durophagous taxa.

## DISCUSSION

Brachyuran ecology has been strongly influenced by changes in functional morphology and biomechanics (34). Cheliped morphology, in particular, has been used to understand durophagous behaviours and prey interactions (25, 33, 37–39) and fiddler crab heterochely (17, 40, 41). Cheliped diversity has traditionally been linked to a broad diversity in prey items (26, 34). However, our integrated assessment of cheliped morphology and ecology demonstrates that these structures record a complex interplay between prey interactions, mechanical effectiveness, environment, and mate attraction specializations (15, 17, 19). This is especially evident in the overlap of traits in morphospace, supporting the many-to-one functional condition for chelipeds and demonstrating the many selective pressures acting on these structures (42, 43).

Cheliped morphospace is defined by finger length and manus morphologies. Slender forms are consistently separated from taxa with shorter, rounded shapes. Some species show considerable heterochely (e.g., ocypodid crabs, Figure 1), and many analyzed species have a robust “crusher” cheliped (often right) and a more slender “cutter” cheliped (often left) (e.g., grapsoids, see Figure 1) (11, 18, 19, 21, 37, 43). Despite this, variation in cheliped morphology between species is much greater than within-species variation (Figure 1). As a result, the distribution of taxa in morphospace is similar regardless of whether right, left, or both claws are included in the analysis (compare Figures 2A, 3A, and 4A). This pattern is visually reinforced by color-coding taxa by ecological trait. Aquatic species show high diversity in cheliped shape whereas terrestrial species are restricted to the lower central part of the morphospace regardless of which claws are analysed (compare Figures 2E, 3E, and 4E). This suggests that cheliped shapes reflect tighter constraints associated with their biomechanical roles in supporting activities in a terrestrial environment.

Although there is overlap across diet categories, there is a signal within the functional landscape (Figures 2B, 3B, 4B) that reflects a distinction in feeding mechanics. Elongated fingers are used to grasp and manipulate softer prey, while more robust chelipeds with larger denticles are associated with durophagous feeding (20, 21, 37). This distinction is reinforced when the functional landscape and diet information are considered together (Figures 2B, D, 3B, D, 4B, D). Peaks in NRFs within positive PCO1 space occur where durophagous forms are located, while generalists and soft-bodied feeders are associated with troughs in the NRF landscape. Durophagous forms differ significantly from other species in NRF (Mann-Whitney test = 1235, *p* < 0.0001), even if those coded as “both” are lumped with completely durophagous species (M-W test = 1181, *p* = 0.002). These functional optima and suboptima demonstrate trade-offs between morphologies optimised for force output (more robust forms) and those that have application within manipulation, or male appendage hypertrophy for sexual display (11, 18, 19, 21, 37, 43). Further, inconsistent distribution of higher NRFs across the functional landscape demonstrate how mechanical efficiency, finger length, and manus shape interplay across appendage morphologies.

Another avenue of considering the NRF data is within the context of cheliped reach and manipulative capacity. Taxa with elongated fingers occupy -PCO1 space, coinciding with troughs in the NRF landscape. These forms likely sacrifice force transmission for greater reach, enabling interception and manipulation of soft-bodied prey (13). Conversely, durophagous taxa occupy +PCO1 space where NRF peaks are highest. Here, shorter and more robust finger morphologies maximise force output at the cost of reduced reach (13, 44). Chelipeds used in male visual display occupy a distinct functional region, in -PCO2 space.

These appendages are characterised by hypertrophy and enlarged muscle masses associated with combat, rather than geometric optimisation for force or reach (30, 40, 45). Stridulatory forms cluster in regions of intermediate to high NRF. Their manus geometry has been optimised for sound production requirements, distinct from prey capture (13). Taken together, the functional landscape of brachyuran chelipeds is structured along intersecting axes of force output and reach. The ecological and behavioural diversity of the group reflects a complex interaction between these demands and evidences different selection pressures on the same morphology.

Despite the substantial differences in NRFs, biomechanical models surprisingly demonstrate that all chelipeds analyzed function similarly, as they all show similar stress distributions. The FEMs therefore evince the mechanically simple function that contrasts with the complex functional landscape derived from NRFs (44). Further, despite modifications across the group, chelipeds are examples of homogenous structures (11, 19).

The overlap of ecologically and phylogenetically disparate taxa across shared regions of morphospace provides strong support for many-to-one mapping of cheliped form to function (46). Many-to-one mapping predicts that similar mechanical performances can be achieved by a range of distinct morphologies, effectively buffering taxa against the fitness consequences of morphological variation (47, 48). This relaxes stabilising selection on shape (47, 48). This is evidenced here with a broad distribution of generalist and soft-diet feeders across the functional landscape that share comparable NRF values, despite distinct morphospace positions. The absence of phylogenetic signal in cheliped shape further supports this interpretation. Furthermore, the morphological diversity observed within superfamilies is indicative of drift within broad performance plateaus. Conversely, stronger selective pressures are reserved for the functional extremes of the landscape, which has resulted in highly robust durophagous, or hypertrophied display morphologies. That phylogenetic signal in NRF values is much weaker in right claws, which are typically those that are more robust or hypertrophied in heterochronic species, also supports this interpretation.

The optima of NRFs are mostly clustered about the highest diversity of taxa. Chelipeds capable of producing high forces overall have comparable morphologies. One major difference is peak NRFs for hypertrophied male fiddler crab (Ocypodoidea) chelipeds. These appendages are used in display and fighting other males (18, 27) and have large muscle masses capable of outputting high forces, associated with combat (40). The higher NRFs determined here therefore represents evidence for biomechanical optimisation associated with combat (49).

Modelled fossils are located proximal to generalist feeders within the cluster of models in morphospace analyses. The fossil forms show a more stout and robust morphology that may represent ancestral conditions. This has appeared to have been maintained within Grapsoidea, Pseudothelphusoidea, and Xanthoidea. Chelipeds that may have been used in durophagy arose in the late Mesozoic (50–52). While it has been hypothesized that the origin of durophagous brachyurans drove a Mesozoic arms race with biomineralized molluscs (25, 53), the data herein support potential facultative durophagy in fossil crabs. Subsequent shifts towards life modes involving derived, sexual display-like characters were likely modified from a body-plan initially optimised for predation (53). Brachyuran chelipeds therefore do not appear to have been subject to strong developmental constraints (40).

Aquatic taxa range across the morphospace, reflecting a broad morphological flexibility for life in marine and freshwater environments. Exclusively terrestrial, air-breathing forms are well differentiated within the morphospace, showcasing morphological shifts associated with occupation of terrestrial conditions (41, 54). Strikingly, these forms also have exclusively soft diets and show strong selection for male visual display, which is especially evident within Ocypodoidea (18, 21). Moving beyond the ancestral marine conditions has clearly resulted in an array of derived morphologies (14).

It is especially noteworthy that there is almost no phylogenetic structure across the morphospace. Although some groups are moderately distinct (e.g., Majoidea, Ocypodoidea, and Portunoidea), taxa in most superfamilies cluster together in one area of the morphospace. This is substantiated by the *Kmult* statistic that shows no phylogenetic signal in right or left claw shape. Ancestral phylogenetic relatedness is therefore a poor differentiator of cheliped morphology. The overlap of distantly related taxa in the morphospaces presents insight into the evolutionary history of these structures. There is strong evidence for the conservation of potential ancestral morphologies represented by the extinct taxa sampled, as the fossil forms also cluster in this space.

Our examination presented here illustrates quantitatively that brachyuran chelipeds are morphologically diverse structures that have been shaped by feeding ecology, environmental transition, and behavioural specialisation. By integrating 3D GM, FEMs, and functional landscapes, we uncovered that cheliped form is reflective of a morphospace explored by distantly related lineages under different ecological pressures. The absence of phylogenetic signal, combined with evidence for many-to-one mapping, suggests that selection acts most strongly at the functional extremes within the landscape. The observation that fossil forms cluster within regions occupied by extant generalists and durophages indicates that modern durophagous, manipulative, and display-specialised forms arose from a morphologically flexible foundation. Taken together, these findings showcase that cheliped diversity is indicative of functional trade-offs across Brachyura.

### Future directions

Here, we aimed to integrate multiple tools to reconstruct brachyuran cheliped shapes, biomechanics, and evolution. In doing so, projects well beyond the scope of this work arose, and we highlight these as key future directions for further advancing this field.

1. <u>Exploring biomechanical performance of appendages during ontogeny.</u> As male fiddler crabs develop, the manus size decreases relative to overall length of the appendage, which is thought impact the effectiveness of the cheliped (28). Understanding this pattern will provide further insight into the “beautiful weapon” displayed by these forms (28, 41, 55). 3D FEA and functional landscapes represent ideal tools to test if such effectiveness decreases during development. Combining these data within phylogenetic contexts will allow generalisations to be made across Ocypodoidea (40, 45).
2. <u>Understanding the trade-off between speed and strength.</u> Fast closing chelipeds are not strong and strong chelipeds cannot be fast (33, 43). This has not been examined across Brachyura, which reflects a pervasive issue—a lack of large-scale functional morphology and mechanical advantage studies (11). By using 3D data comparable to that collated here, mechanical advantage and indices of closing speed can be derived and examined at scale to the supposed speed-strength trade-off (27, 45).
3. <u>Understanding the relationship between heterochely and diet.</u> A lack of heterochely has been attributed to either not having a predatory life-mode or specialised prey handling (Davie et al., 2015). We have modelled species that lack heterochely (e.g., *Corystes*, *Chionoecetes*, and *Percnon*). However, these genera are recorded as having consumed softer prey (56–58), with FEMs illustrating a similar functionality to other chelipeds. A lack of heterochely thus does not necessitate a lack of predatory life mode, but does support a less durophagous ecology.

## MATERIALS AND METHODS

### Raw 3D data and core dataset

***Specimens*—**Modern brachyuran specimens were selected from the American Museum of Natural History (AMNH) decapod collection. Forms were selected to cover an array of appendage morphologies across Brachyura. Three fossil forms were selected from the University of Alabama Museums Research & Collections (UAMRC) paleontology collection and the AMNH Invertebrate Paleontology collection and analyzed to reconstruct extinct forms within the larger framework (see Supplemental Table 1 for specimen numbers).

***Scan data*—**Specimens were scanned under optimised conditions using a GE-Phoenix v|tome|xs micro-CT scanner and the ‘‘Direct’ tube in the Microscopy and Imaging Facility (MIF) at the AMNH (Supplemental Table 1).

***Segmentation***—Scans were imported into *Mimics* (v. 26.0) and segmented using the ‘Segmenting’ tool. The propodus (manus and pollex), dactylus (terms from Davie et al. (13), Spani et al. (17), and closing muscles for left and right appendages of all specimens were segmented (Supplemental Figure 1). For fossils, the matrix infilling the propodus was segmented as a proxy for muscles, as muscle data are exceptionally rare within the fossil record (59). 3D models of exoskeletal parts and muscles were exported as .stl files from *Mimics* and imported into *Geomagic Studio* (v.2014.3.0). Reconstructions were smoothed in *Geomagic Studio* and the angle between the propodus and dactylus was set to 20–30° [biologically realistic gape values across the group (27, 33)] for biomechanical analyses (Supplemental Table 2).

***Mesh alignment***—Prior to morphometric analyses, dactylus sections were rotated to an occluded state in *Geomagic Studio* to reduce model variation. They were then converted to .ply files in *MeshLab* (v. 2025.07) (60). Meshes were aligned for right-handed claws and left-handed claws separately. Meshes for right-handed claws were also transformed by flipping meshes along the *x*-axis (i.e. creating a “mirror image”). This resulted in a framework in which all meshes could be aligned in one analysis. The transformation was applied in *MeshLab*.

### Biomechanical analyses

***Muscle force calculation*—**Input muscle force is needed for biomechanical models. Muscle cross-sectional area was used to calculate input muscle force (43). Maximum cross-sectional muscle area (CSMA) of segmented muscles was derived from the 3D muscle reconstruction using the ‘Calculate area’ function in *Geomagic Studio*. After CMSA was derived, input force was derived using

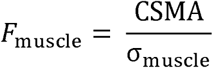

where *F* is input force (N) and σ is muscle stress (Nmm^-2^) (61). We used a σ value of 0.270 Nmm^-2^, a reasonable stress value for cheliped muscular (43). After *F*_muscle_ was calculated, the value was divided by 3 for subsequent biomechanical analyses and muscle loading (Supplemental Table 2)

***Finite element modelling*—**3D biomechanical modelling provides a powerful, non-destructive approach for predicting mechanical performance in animals. In particular, 3D finite element analysis (FEA) can qualitatively and quantitatively estimate how stresses distribute throughout biological structures (62–65). The left and right propodus and dactylus .stl files exported from *Geomagic Studio* were imported as unique sections into *GMSH* (v. 4.15.1) to be solid-meshed as distinct solid homogeneous structures in tet-4 elements (66). Sections were meshed as unique objects and exported as Nastran files for import into *Strand7* (v. 3.0) FEA software. Material properties used for analyses are a Young’s modulus of 15 GPa—a representative value, as decapod moduli range between 1–59 GPa (12, 67, 68) and a Poisson’s ratio of 0.3 (69). Muscle origins and insertions were tessellated as beam elements onto the Nastran models, following applications on scorpion pedipalps (69). Muscle forces (Supplemental Table 2) were assigned to three trusses directed from origins to insertions on the dactylus (49). Three denticles along both the pollex and dactyl located at the proximal, medial, and distal regions were constrained in all directions at their most apical node following Bicknell*, et al.* (69). A hinge between the propodus and dactylus was constructed using links on both appendage sides (69). Colour-coded von Mises (VM) stress maps were generated after solving models. Maps all were scaled to a maximum value of 20 MPa for comparison (Supplemental Table 2; Figure 1). This value was selected as it effectively showcased stress distributions across all models. Loaded and solved *Strand7* models are presented as Supplemental Dataset 1 found at https://figshare.com/s/0a2e81a7877663d47334. This pipeline is comparable to other studies that applied FEA to chelipeds (12, 70). We followed the Zhang*, et al.* (12) approach of using single material properties and did not separate the outer most cuticle layer (70) to allow easier comparisons between models.

***Node reaction forces***—Node reaction forces (NRFs) are force output values from constrained nodes in the finite element models (FEMs). These values calculate how force input is translated into realised output. All constrained nodes were examined, and force values from the six triangles around the nodes were derived and averaged to determine output forces at constrained nodes (Supplemental Table 3). The average values per node were then averaged again to produce a mean NRF for left and right appendage models. This resulted in a dataset in which appendages could be explored independently, or relative to each other.

### Three-dimensional geometric morphometrics (3D GM)

***Alignment****—*Meshes were aligned using generalized Procrustes surface alignment in the software *GPSA* (v. 20160308) (https://morphlab.sc.fsu.edu/software/gpsa/index.html). This alignment approach uses the Iterative Closest Point algorithm to associate each point on one surface with its nearest neighbor on another surface. This calculates and applies a transformation to minimize a cost function based on the distances between the point pairs. These steps are repeated until the distance cannot be reduced further. The cost function is a modification of the Procrustes distance that allows application to surfaces with different numbers of vertices (71). Following alignment, we ran a principal coordinates analysis of the distance matrix of pairwise Procrustes surface metrics, as implemented in the *GPSA* software. The ordination scores (PCO) were then used to visualize variation in cheliped shape (Supplementary Tables 5–7).

***Sensitivity to resolution of mesh****—*To determine if the number and distribution of vertices/faces across models of different shapes and sizes impacted alignment, a subset of seven models was selected and subsampled at resolutions of 50,000; 30,000; 20,000; and 10,000. This subset was aligned against the same reference and heat maps of the variance from the mean shape were compared to each other. The same patterns of variation across cheliped shape were observed regardless of model resolution (Supplemental Figure 2). For all subsequent analyses, all models therefore were sampled at 50,000 faces within *Geomagic Studio.* In practice, the analytical software frequently yields small variations of face numbers in sampled models (e.g., the *Ovalipes ocellatus* right cheliped has 49,996 faces, i.e., slightly fewer than 50,000).

***Selection of prototype (reference shape for alignment)****—*Alignment is applied to pairs of surfaces. As such, alignment is sensitive to reference shape (prototype) selection, and the superimposition of one surface onto another will not necessarily produce the same superimposition if reversed (71). The best prototype was identified through trial and error, starting with shapes considered close to the mean shape, and assessing results by visually comparing aligned shapes in *Meshlab*. The most common misalignment was 180° in either horizontal or vertical directions, such that either the manus and fingers, or pollex and dactylus were misaligned. The right cheliped of *Ocypode urvillii* was the first identified reference shape against which no chelipeds were misaligned, and was used as the prototype for all analyses.

### Phylogenetic analyses

***Molecular data—***A dataset of 10 Sanger sequenced genes was constructed as shown by Wolfe*, et al.* (14), to resolve brachyuran relationships (Supplemental Dataset 2). These were used instead of a phylogenomic approach as genomic scale data are unavailable for many scanned taxa. Genes from an exact species match, congener, or relative in the same subfamily were used (Supplemental Table 4). For five taxa (*Acanthothelphusa congoensis*, *Libinia dubia*, *Libinia emarginata*, *Ocypode cursor*, *Paradasygyius*L*depressus*), no accurate molecular data for a close enough relative were identified. These were therefore excluded from the phylogeny and ‘float’ in phylomorphospace. Molecular data (for relatives) were identified to place the fossil specimens of *Chaceon peruvianus* (within the extant family Geryonidae) and *Pulalius vulgaris* (within the extant superfamily Carpilioidea; Schweitzer et al. (72)). As a sensitivity assessment for the potential impacts of this approach, analyses including and excluding these two terminals were presented (Supplemental Figures 4, 5).

Substitution was not possible for *Dakoticancer australis* as it falls within a completely extinct podotreme group. Data for the 10 genes, nuclear (18S, 28S) and mitochondrial (12S, 16S) ribosomal RNA (rRNA) genes, and six nuclear protein-coding genes [AK (arginine kinase), enolase (phosphopyruvate hydratase), GAPDH (glyceraldehyde 3-phosphate dehydrogenase), H3 (histone 3), NaK (sodium–potassium ATPase α-subunit), and PEPCK (phosphoenolpyruvate carboxykinase)] were downloaded from the National Center for Biotechnology Information (NCBI). Sequences (accession numbers in Supplementary Table 4) were identified either by querying the nucleotide records by taxon, or downloading the annotated sequence from full mitogenomes. All downloaded sequences were queried (73) against their labeled taxon using *BLASTn* (for both rRNA and protein-coding genes) to prevent inclusion of incorrectly identified sequences. A minimum of two genes were required for each extant taxon to be included in the analysis, with an average of five genes per tip taxon.

***Phylogenetic analyses—***rRNA and protein-coding genes were individually aligned in *MAFFT* (v.7.31) (74) using the L-INS-i algorithm. Protein-coding genes were translated to check for pseudogenes. Gene alignments were masked in *GBlocks* (v.0.91b) (75) under “less stringent” parameters to remove regions with poor homology. Masked alignments were concatenated with *phyx* (76), resulting in 6983 nucleotide characters.

Protein-coding genes were partitioned by codon position *a priori*. Following that, best-fitting partitions were selected by an edge-linked proportional model (77) and best-fitting substitution models with *ModelFinder* (78) in *IQ-TREE* (v.1.612) (79). The merged scheme was used to estimate the concatenated maximum likelihood (ML) phylogeny, also in *IQ-TREE*, using 10 independent runs with perturbation strength 0.2 and stopping at 500 iterations, selecting the topology with the highest log likelihood. Ultrafast bootstrap values were calculated from 1000 replicates using the -bnni flag (80).

Phylogenetic analyses using Bayesian inference (BI) were conducted in *MrBayes* (v.3.2.7) (81). The analysis implemented four runs of four chains each (for 35 million generations), with 25% burnin. Convergence required reaching standard deviations of split frequencies <0.01, reaching effective sample size >200 for every parameter, and overlapping posterior distributions in *Tracer* (v.1.7.1) (82).

### Phylomorphospace

Combining the ML tree that included fossils with PCO data allows construction of phylomorphospaces for left and right chelipeds, as well as both appendages in the same space. The phylogenetic topology determined from the phylogenetic analyses was imported into *R* (v. 4.5.2) using *ape* (83, 84) and matched to the 3D GM dataset (Supplemental Tables 5–7). Species excluded from the ML terminals and those not included in the 3D GM dataset were pruned to match the tree topology and PCO datasets and then rooted with *Dromia erythropus* at the base (14). The positions of internal nodes within PCO space were estimated using maximum likelihood, as implemented in the *phylomorphospace* function in the *phytools* R package (85). Individual terminals were overlaid as points onto PCO1 and PCO2 bivariate spaces. Any species modelled using 3D GM data that were pruned were presented as floating. Both terminals and branches were colour-coded by superfamily to highlight higher-level taxonomic structure.

### Functional landscape

A functional landscape illustrates a performance property superimposed as a third-dimension onto a bivariate morphospace (86). This differs from adaptive landscapes because only one metric is considered and neither fitness characteristics (87, 88) nor natural selection are explored (89–93). NRFs were used to construct a functional landscape to contextualise the FEA outputs with the morphospace. To do so, NRFs were log_10_ normalised (log_10_N) (Supplemental Table 3) and values were interpolated across the PCO1 and PCO2 space using the *akima R* package (84, 94). Interpolation was gridded (150×150) and rendered as a colour-scaled performance space.

### Phylogenetic signal

Phylogenetic signal in NRF was assessed using the *K* statistic (95) as implemented in the *phytools* R package (85). Phylogenetic signal in cheliped shape was assessed using *Kmult*, a generalization of the K statistic for highly dimensional multivariate data (96). *Kmult* was calculated for the full set of PCO scores. For both phylogenetic signal analyses, *p*-values were based on 1000 randomizations.

### Ecological traits

Cheliped multifunctionality reflects the evolutionary outcome of an array of underlying selection processes over time. To explore this, ecological trait data were collected from natural history literature (Supplemental Tables 8; 9). For coding, the trait needed to be observed for a given species or its sister species or reported to be conserved across an entire clade. When this information was unavailable, the trait was coded as ‘NA’.

***Diet—***Brachyuran diets were classified as hard foods, soft foods, or both. ‘Hard’ foods, indicative of durophagy, represent biomineralized organisms, such as shelled molluscs, sea urchins, or other crabs (25). ‘Soft’ foods represent everything else, including vertebrates, as brachyurans use chelipeds to remove flesh and employ mandibles to masticate smaller individuals. Species that consume both hard and soft food were classified as ‘both’. Ideally, gut content analysis (97) would more fully inform these categorizations. However, as those studies are rare, behavioral observations were mostly used to assign diet categories (Supplemental Tables 8; 9). Critically, a complete assessment of a species’ diet across its range, seasonality, and other factors are rarely, if ever, available.

***Environmental habitus medium—***Occupation of distinct environments may have influenced the overall morphology and the morphofunctionality of chelipeds. As such, whether a species is mostly submerged in water or lives within terrestrial settings also was evaluated. Species inhabiting upper intertidal zones, mangrove habitats, or with diurnal cycles between aquatic and terrestrial were characterised as ‘both’ for this variable.

***Courtship signals—*** In addition to ecological traits, sexual display via courtship signals also were considered, because visual signaling by claw waving has been correlated with biomechanical function (41, 54). Brachyuran sexual display is best documented in male fiddler crabs (27, 28, 40, 41). Stridulation—sound production—is another sexual signal found across multiple brachyuran families (98). For these data, stridulatory behaviors associated with cheliped use were included. Notably, cheliped stridulation is most strongly expressed in male fiddler crabs and likely influenced their morphological evolution (41, 99).

## Supporting information

Supplemental Dataset 2. Complete dataset used for phylogenetic analysis.

Supplemental Dataset 2. Complete dataset used for phylogenetic analysis.

Supplemental Figure 3. Full phylogenetic hypothesis for Brachyura based on the topology from the ML concatenated analysis

Supplemental Figure 4. Phylogenetic hypothesis for Brachyura based on the topology from the ML concatenated analysis, but with the two fossils (Chaceo

Supplemental Figure 5. Full phylogenetic hypothesis for Brachyura based on the topology from the Bayesian concatenated analysis.

## ACKNOWLEDGEMENTS

This research was supported by an Australian Research Council grant DE250100256 (to R.D.C.B.) and by an AMNH-RGGS MAT Program Postdoctoral Fellowship (to R.D.C.B.), and by the US National Science Foundation DEB #1856679 (to J.M.W.). We thank Lily Berniker, Hilary Ketchum, and Peter Daly for help with the AMNH collections. We thank Lisa Nink for feedback on the manuscript.

---

Supplemental Figure 1. Simplified depiction of cheliped showing major parts used in 3D FEMs.

Supplemental Figure 2. Sensitivity of mesh resolution on shape alignment. Left column shows mesh for prototype specimen (here the left claw of *Sudononautes aubryi*) comprised of 10,000, 20,000, 30,000, and 50,000 faces. Right column shows heat maps of the shape variance across seven specimens sampled at the respective resolutions and aligned using the prototype. The heat map is projected onto the estimated mean shape. Blue indicates low values, red indicates high values. These values are calculated from the covariance matrix of the nearest neighbor points for each point on the prototype. Heat maps are very similar across all analyses.

Supplemental Figure 3. Full phylogenetic hypothesis for Brachyura based on the topology from the ML concatenated analysis. Taxon labels follow the spellings from NCBI and do not directly correspond to scanned species. Values at nodes represent ultrafast bootstraps. Branches colored by superfamily and labelled by family.

Supplemental Figure 4. Phylogenetic hypothesis for Brachyura based on the topology from the ML concatenated analysis, but with the two fossils (*Chaceon peruvianus* and *Pulalius vulgaris*; substituted by molecular data from *Chaceon* sp. and *Carpilius maculatus*, respectively) omitted. Taxon labels follow the spellings from NCBI and do not directly correspond to scanned species. Values at nodes represent ultrafast bootstraps.

Supplemental Figure 5. Full phylogenetic hypothesis for Brachyura based on the topology from the Bayesian concatenated analysis. Taxon labels follow the spellings from NCBI and do not directly correspond to scanned species. Values at nodes represent posterior probabilities.

**Supplemental Table 1.** Meta-data associated with modelled specimens, including scan conditions.

**Supplemental Table 2.** Data associated with 3D biomechanical analyses. Includes input force data, node reaction forces and mean von Mises stress values.

**Supplemental Table 3**. Raw data for bricks about constrained nodes used to average node reaction forces.

**Supplemental Table 4.** Sample information for molecular data, with corresponding taxonomic substitutions where necessary. Dark greyed out taxa are not included in the phylogeny. Light greyed out taxa are included but represent fossils.

**Supplemental Table 5.** Data used to construct phylomorphospaces for left cheliped. Includes averaged von Mises stress values and PCO scores.

**Supplemental Table 6.** Data used to construct phylomorphospaces for right cheliped. Includes averaged von Mises stress values and PCO scores.

**Supplemental Table 7.** Data used to construct phylomorphospaces for both appendages in the same space. Includes averaged von Mises stress values and PCO scores.

**Supplemental Table 8.** Trait data used to contextualize morphospaces.

**Supplemental Table 9.** References associated with trait codings in Supplemental Table 8.

**Supplementary Code 1**. R code associated with presented figures and analyses.

**Supplemental Dataset 1.** Complete dataset of biomechanical models and .stl files of modelled chelipeds. https://figshare.com/s/0a2e81a7877663d47334

**Supplemental Dataset 2**. Complete dataset used for phylogenetic analysis.

