## Supplemental Dataset 2. Complete dataset used for phylogenetic analysis. for "Complex interplay of biomechanics and ecology influenced crab claw morphology evolution": README.txt

This is the README file for all supplementary data associated with: 

***************Complex interplay of biomechanics and ecology influenced evolution of crab claw morphology***************

Each folder and its file contents are described below. 

Fasta (.fasta) is a file type for sequence data and multiple sequence alignments.
Nexus (.nex) is a common file type for phylogenetic data and multiple sequence alignments, and the required runfile to use MrBayes for phylogenetic analysis.
Newick (.tre, .trees, .treefile) files are phylogenetic tree and chronogram outputs from multiple programs, which can be visualized directly e.g., with Figtree.

***

1_raw_gene_alignments (folder)
Contains 10 .fasta files for the multiple sequence alignment of each gene in MAFFT, before any processing (i.e., as downloaded from NCBI). Also contains 10 .fasta files that have been aligned in MAFFT.

2_processed_gene_alignments (folder)
Contains 10 .fasta files for the multiple sequence alignment of each gene in MAFFT, which have first been processed with GBlocks. 

3_concatenated_ML_analysis (folder)
Contains the concatenated multiple sequence alignment (based on the processed gene alignments in folder 2), model and partitions schemes from IQ-TREE, and the resulting concatenated gene tree. 
3_concatenated_tree.contree - is visualized in full in Fig S3.
5_renamed_iqtree.contree - is the basis for phylomorphospace analyses.
8_concatenated_tree_nofossil.contree - is visualized in full in Fig S4.

4_concatenated_Bayesian_analysis (folder)
Contains the runfile for MrBayes (includes the concatenated alignment in folder 3), sampled parameter files, and the resulting concatenated Bayesian tree.
2_concatenated_bayes_tree.tre - is visualized in full in Fig S5.
