## Supplementary figures and images for "Complex interplay of biomechanics and ecology influenced crab claw morphology evolution"

### Supplemental Figure 3. Full phylogenetic hypothesis for Brachyura based on the topology from the ML concatenated analysis

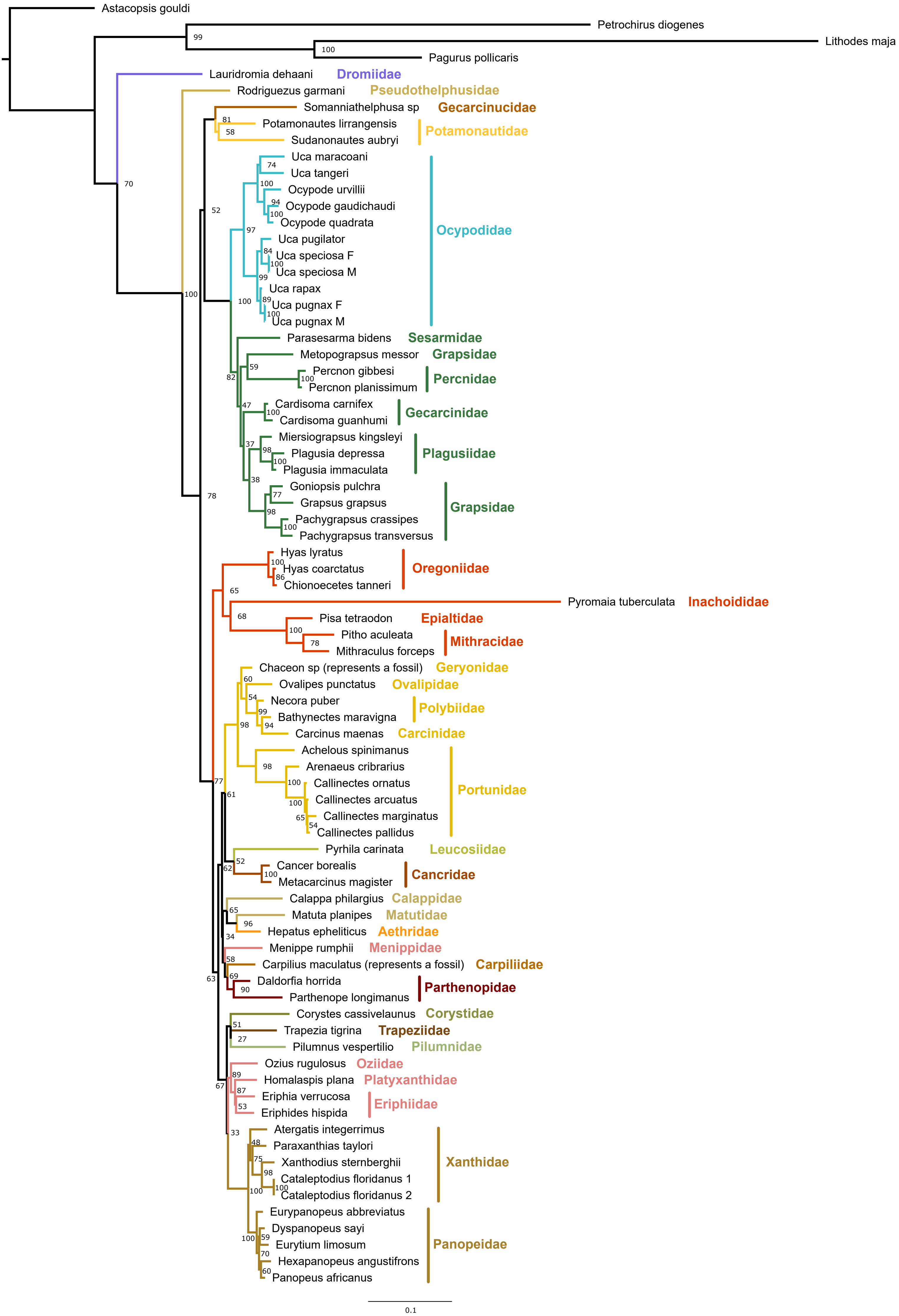

### Supplemental Figure 4. Phylogenetic hypothesis for Brachyura based on the topology from the ML concatenated analysis, but with the two fossils (Chaceo

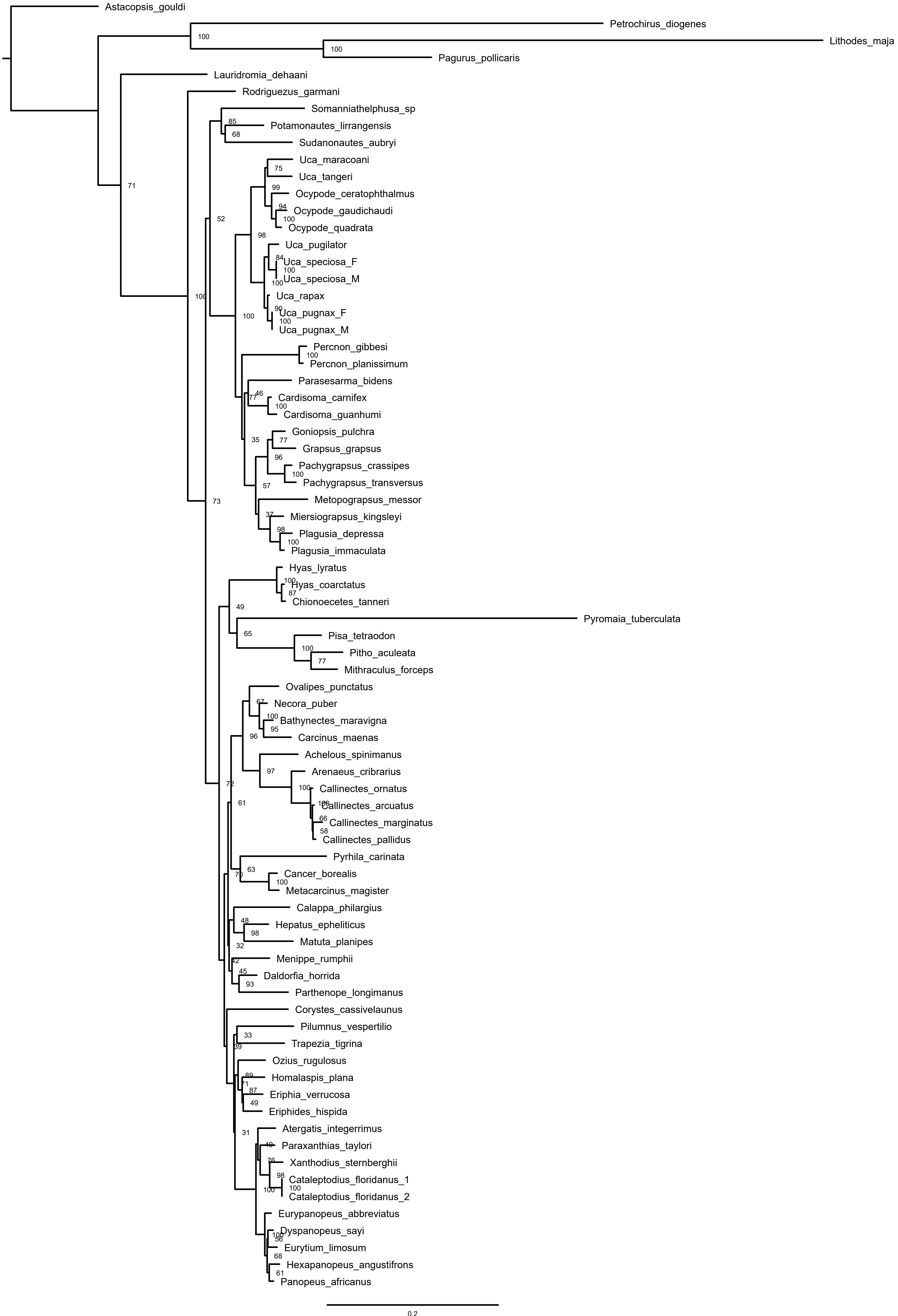

### Supplemental Figure 5. Full phylogenetic hypothesis for Brachyura based on the topology from the Bayesian concatenated analysis.

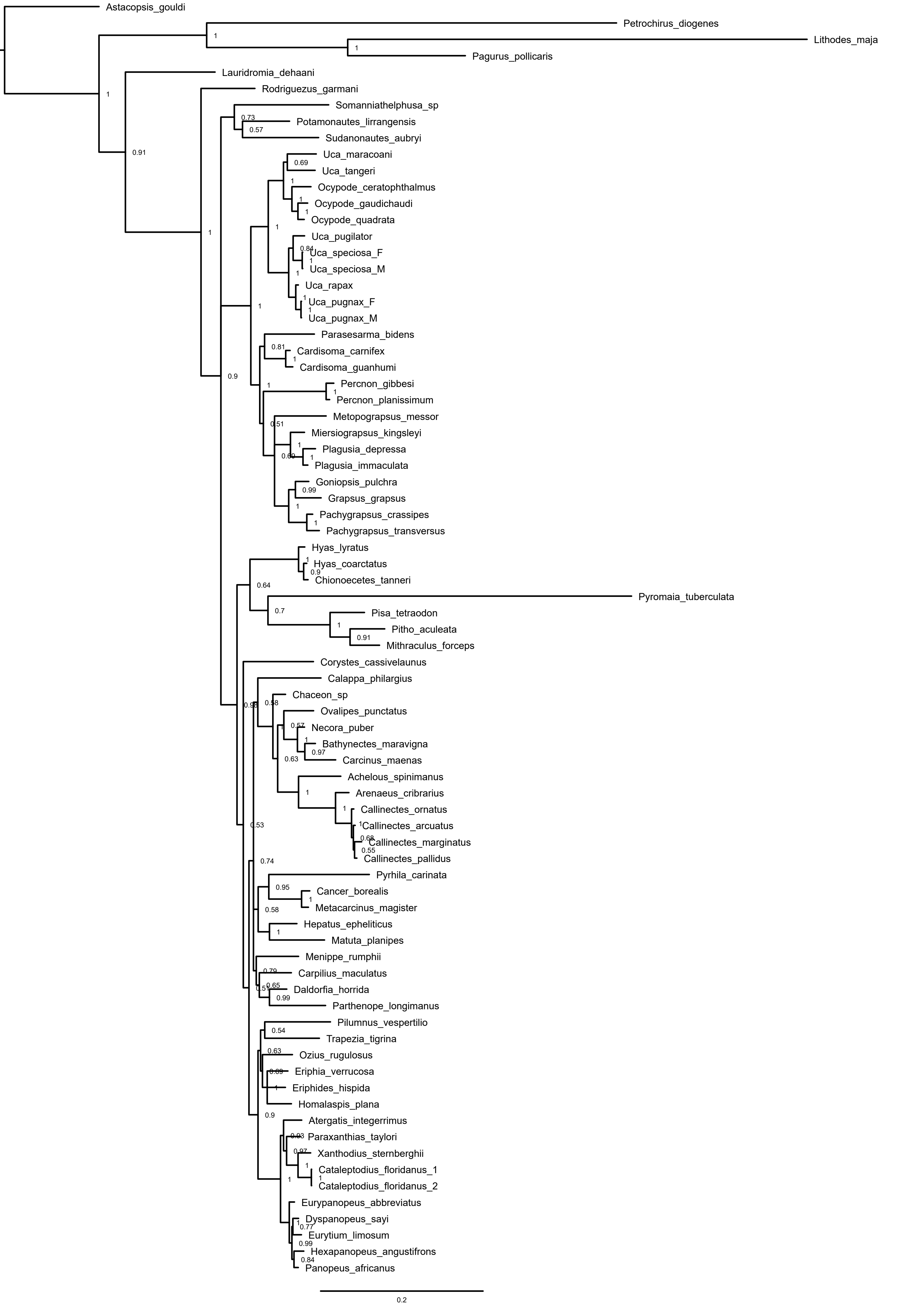
